# The Crystal Structure of Klebsiella pneumoniae FeoA Reveals a Site For Protein-Protein Interactions

**DOI:** 10.1101/514059

**Authors:** Richard O. Linkous, Alexandrea E. Sestok, Aaron T. Smith

**Affiliations:** Department of Chemistry and Biochemistry, University of Maryland, Baltimore County, Baltimore, Maryland, 21250 USA

## Abstract

In order to establish infection, pathogenic bacteria must obtain essential nutrients such as iron. Under acidic and/or anaerobic conditions, most bacteria utilize the Feo system in order to acquire ferrous iron (Fe^2+^) from their host environment. The mechanism of this process, including its regulation, remains poorly understood. In this work, we have determined the crystal structure of FeoA from the nosocomial agent *Klebsiella pneumoniae* (*Kp*FeoA). Our structure reveals an SH3-like domain that mediates interactions between neighboring polypeptides *via* intercalations into a Leu zipper motif. Using docking of a small peptide corresponding to a postulated FeoB partner binding site, we demonstrate the *Kp*FeoA can assume both ‘open’ and ‘closed’ conformations, controlled by peptide binding. We propose a model in which a ‘C-shaped’ clamp along the FeoA surface mediates interactions with its partner protein, FeoB. These findings are the first to demonstrate atomic-level details of FeoA-based protein-protein interactions, which could be exploited for future antibiotic developments.

---

The acquisition of iron is an essential virulence factor for the establishment of infection by a wide array of bacterial pathogens,^1–2^ including one of the major causative agents of nosocomial (hospital-acquired) infections, *Klebsiella pneumoniae*.^3-4^ The environmental source of iron is typically the host, where it may be found in multiple oxidation and coordination states, necessitating pathogens such as *K. pneumoniae* to adapt to acquire iron in ferric (Fe^3+^), ferrous (Fe^2+^), and even chelated forms. Under oxidizing conditions, siderophore- and heme-based acquisition systems are essential to stabilize, to solubilize, and to transport ferric iron.^1^ However, under acidic, micro-aerobic, and/or anaerobic conditions, such as those found in the gut or within biofilms, iron may be prevalent in the reduced, ferrous form.^5–6^ Because ferrous iron has differences in solubility, lability, and even coordination properties compared to ferric iron, bacteria such as *K. pneumoniae* must employ orthogonal transport systems to acquire and to handle Fe^2+^.^5–6^.

The most prevalent prokaryotic transport system dedicated to the transport of Fe^2+^ is the <u>fe</u>rr<u>o</u>us iron transport system, also known as Feo (Fig. 1).^6^ In *K. pneumoniae,*this system is found along the *feo* operon (Fig. 1A), which encodes for three proteins: FeoA, FeoB, and FeoC.^7^ FeoA and FeoC are predicted to be small (~8 kDa), cytosolic proteins, whereas FeoB is a large (~90 kDa), polytopic membrane protein bearing a N-terminal GTP-binding domain (NFeoB; Fig. 1B). These three proteins are postulated to function in concert to regulate the movement of ferrous iron from the periplasm into the cytosol (Fig. 1B), where it is presumably handed off to an unknown ferrous iron chaperone for assembly into iron-containing proteins and/or intracellular storage.^6^

**Figure 1.**
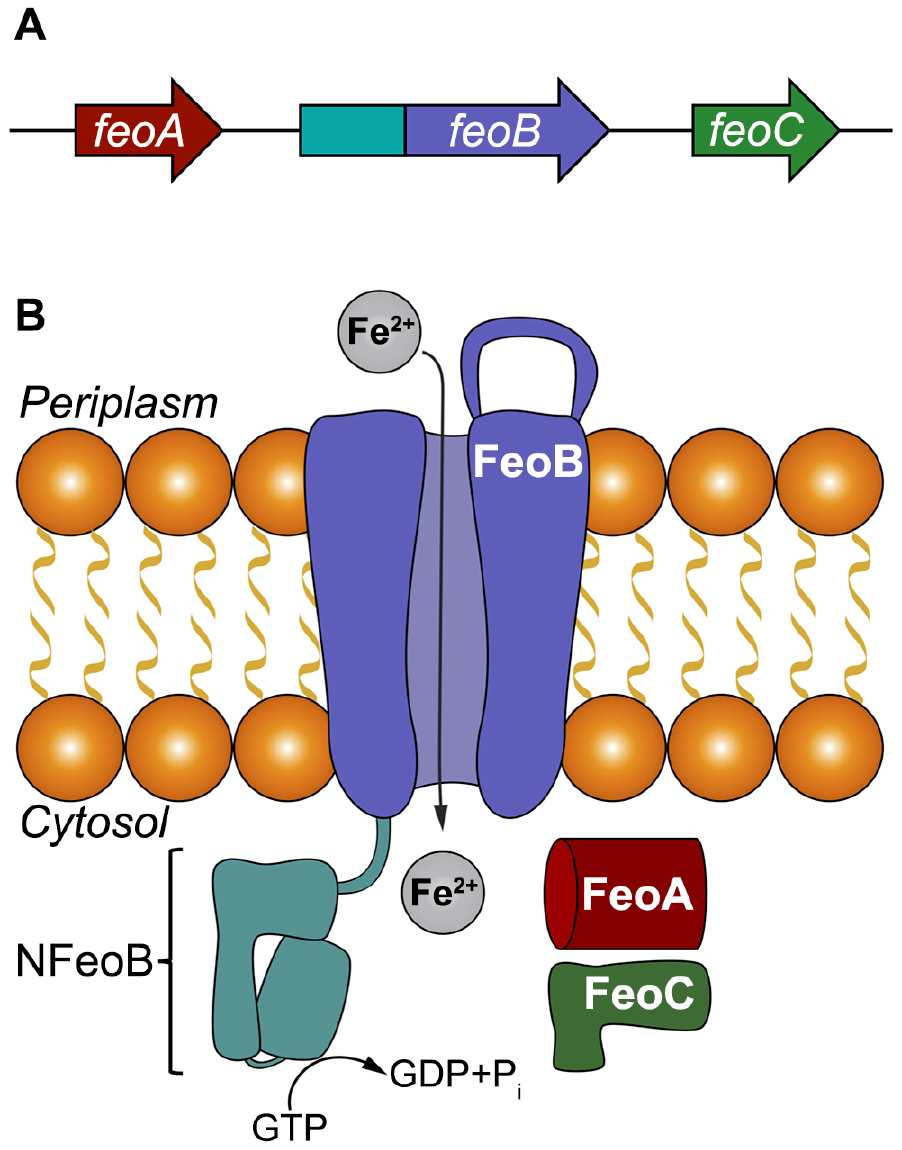
The Feo system in *K. pneumoniae.* **A**. The *feo* operon within *K. pneumoniae* encodes for three proteins: FeoA, FeoB, and FeoC. **B**. In *K. pneumonia*, FeoA (red) and FeoC (green) are small, cytosolic proteins that are predicted to function as accessories to ferrous (Fe^2+^) transport across the intracellular membrane, mediated by the polytopic membrane protein FeoB (purple). At the N-terminus of FeoB is the soluble GTP-binding domain known as NFeoB (teal), which is capable of hydrolyzing GTP and may drive ferrous iron transport in an active manner.

We recently described the preparation and biophysical properties of detergent-solubilized *K. pneumoniae* FeoB (KpFeoB).^7^ Of note was the ability of KpFeoB to hydrolyze GTP at a rate comparable to the hydrolysis of ATP by some ABC^8–9^ and P^1B^-ATPase^10–11^ metallotransporters, suggesting that ferrous iron transport could be driven in an active manner by GTP hydrolysis. However, this hydrolysis rate (~0.1 s^-1^) is still considered sluggish by means of most active transporters.^7^ The slow rate of GTP hydrolysis by FeoB has lead us and others to consider that an additional stimulatory factor may exist to upregulate hydrolysis under changing intracellular conditions in order to drive ferrous iron uptake.^6–7, 12–14^ It is postulated that this stimulatory factor is FeoA.

Several three-dimensional structures of FeoA exist,^13, 15^ and these structures reveal the presence of an Src-homology 3 (SH3)-like fold. SH3 folds, which are common to eukaryotes and are characterized by small (<100 amino acids) β-barrels, are often involved in mediating protein-protein interactions and are even utilized to activate eukaryotic GTPases.^16–19^ SH3 folds typically interact with binding sites on partner proteins bearing a consensus motif of PxxP, with “x” frequently being a hydrophobic amino acid.^20^ To our knowledge, every FeoB sequence that we have examined contains a candidate PxxP binding site, leading to the consideration that this site is the location for FeoA-FeoB interactions. However, how FeoA may facilitate this interaction remained unknown until now.

In order to investigate the function of *Kp*FeoA, we cloned, expressed, and purified *Kp*FeoA to good yield and high purity (Supplemental Methods; Fig. S1). Circular dichroism studies indicated our overexpressed, purified protein was folded (data not shown), and gel filtration studies demonstrated the presence of two oligomeric species whose molecular weights were consistent with monomeric (~10 kDa) and dimeric (~20 kDa) (His)_6_-tagged *Kp*FeoA (Fig. S2). We observed no trimeric or higher-order oligomerization of *Kp*FeoA under these conditions. Moreover, our oligomeric states appeared to be static under our gel filtration conditions, as dilution of dimeric *Kp*FeoA and reinjection preserved oligomeric homogeneity (Fig. S2). To characterize these states further, we attempted to crystalize both oligomeric species independently; however, we were only successful in generating diffraction-quality crystals with monomeric *Kp*FeoA in the crystallization drop.

Crystals of unmodified, tagged *Kp*FeoA diffracted to <2 Å, and our best native dataset was processed to a resolution of 1.57 Å (Table S1). Despite exhaustive efforts to utilize molecular replacement (MR) to determine phases, including the use of an unpublished NMR structure of *Kp*FeoA, we were unable to find a suitable MR solution, suggesting our structure adopted a conformation distinct from previously determined structures. Subsequently, we expressed, purified, and crystallized SeMet-derived *Kp*FeoA (Supplemental Methods), and were able to establish phases utilizing single-wavelength anomalous dispersion (SAD). This model then was used to establish phases definitively for our native data via MR. Our final refined model converged to an *R*_work_/*R*_free_ of 0.174/0.193 (Table S1), and all residues comprising the full length of the *Kp*FeoA polypeptide (1-75) and part of the tag cleavage site (76-80) were unambiguously present in the electron density (Fig. S3).

Initial inspection of a single monomer comprising the X-ray crystal structure of *Kp*FeoA reveals the expected SH3-like fold present in other FeoA structures (Fig. 2A).^13, 15^ In particular, we observe five β strands (β1-β5) that comprise the β barrel, complemented by two additional *α* helices (*α*1,*α*3) and a helical turn (*α*2) that appear to be unique to FeoAs (Fig. 2B).^13, 15^ A fourth, short helix (*α*4) appears at the C-terminal tail in the visible electron density, but this helix is composed of residues that are part of the tag cleavage site and are not part of the native sequence.

**Figure 2.**
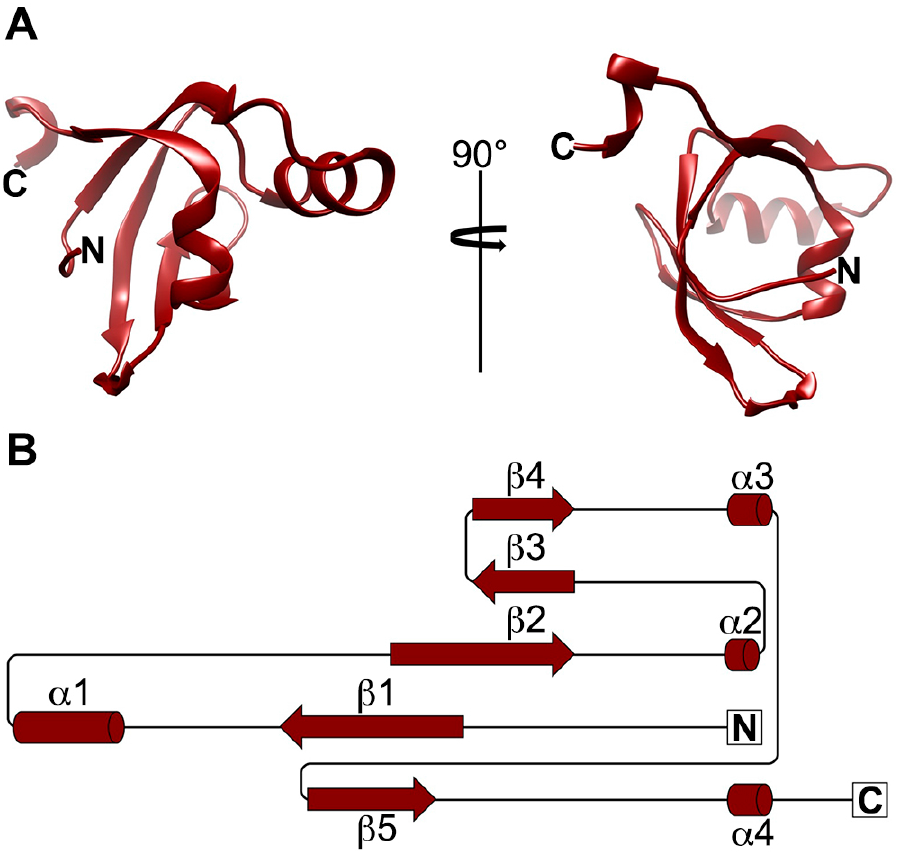
The crystal structure of a single *KpVeoA* polypeptide (PDB ID 6E55). **A**. The ribbon structure of *Kp*FeoA reveals a small ß barrel comprising an SH3-like fold. The right panel represents a 90° rotation of the left panel. *B*. Secondary-structure topology diagram of monomeric *Kp*FeoA. α helices and β sheets are numbered sequentially. “N” and “C” represent the location of the N- and C-termini, respectively.

Intriguingly, analysis of the *Kp*FeoA asymmetric unit (ASU) composed of 6 polypeptides reveals relevant interactions among neighboring *Kp*FeoA molecules (Fig. 3A). Within the ASU there are 2 *Kp*FeoA dimers and 2 *Kp*FeoA monomers. Comprising the dimers, two independent *Kp*FeoA chains participate in hydrophobic intercalations into a Leu zipper motif present on their dimeric partner *Kp*FeoA chains, allowing for the exclusion of a modest amount (~29 Å^2^) of hydrophobic surface area (Fig. 3B,C). The Leu zipper motif on a single *Kp*FeoA polypeptide is composed of four residues along a surface ridge that forms a “C-shaped” clamp (Fig. 3D): Leu^26^ and Leu^29^ (present along α1) and Leu^58^ and Leu^60^ (present along β4). Intercalated into this hydrophobic zipper are two residues of the neighboring *Kp*FeoA monomer along β5: Ala^72^ and Ala^74^ (Fig. 3D). This intercalation appears to displace Leu^29^ from the central portion of the zipper towards the Ala residues along β5 (Fig 3D). This interaction is repeated throughout the crystal (Fig. S4), and thus the two *Kp*FeoA “monomers” without partners in the ASU interact with the β5 of the adjacent ASU, repeating these interactions throughout the entire crystalline lattice.

**Figure 3.**
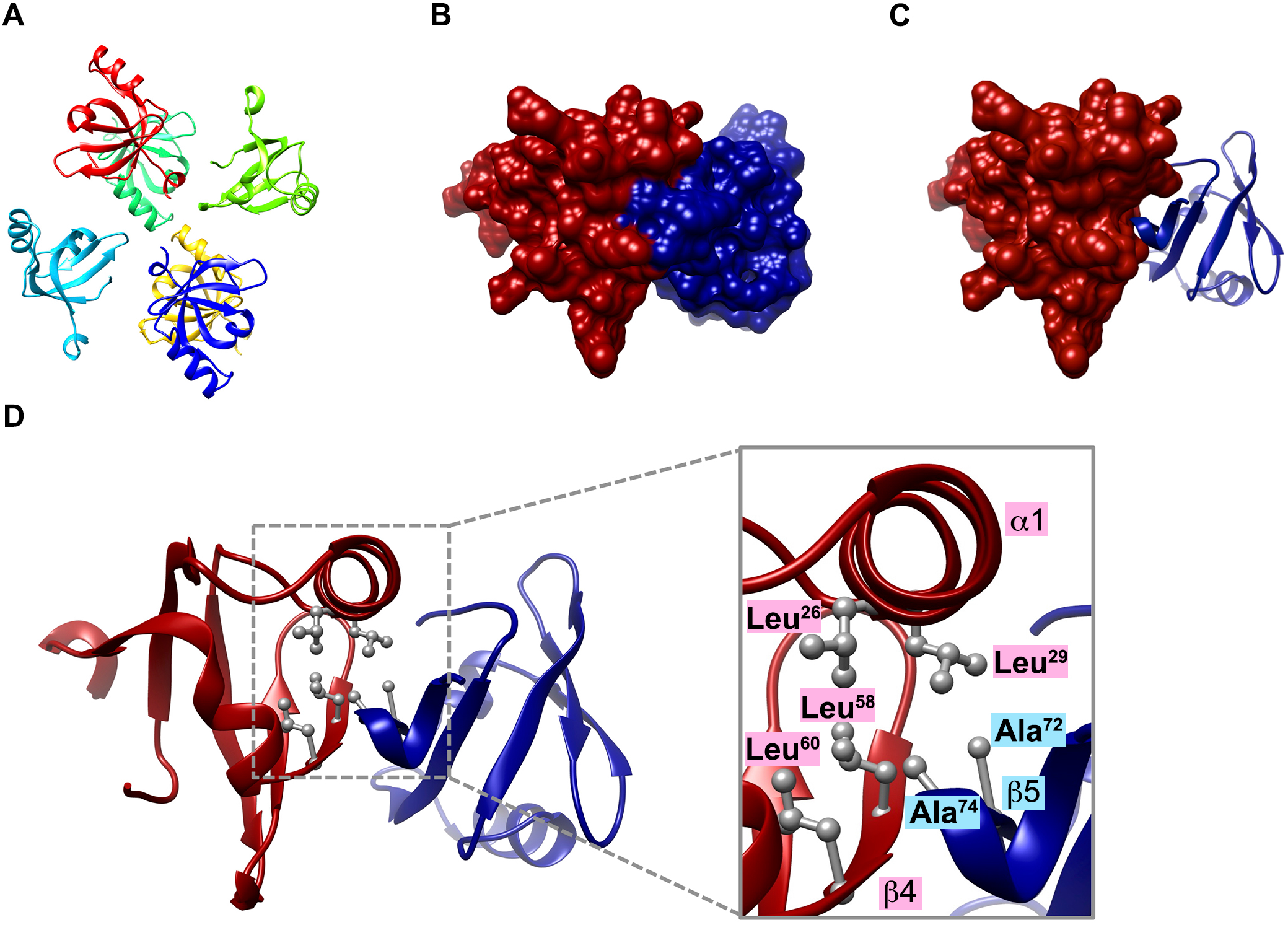
Features of *Kp*FeoA interactions. **A.** Ribbon representation of the asymmetric unit (ASU) of the crystal structure of *Kp*FeoA, which contains 6 molecules. **B.** Space-filling model of two adjacent, interacting *Kp*FeoA polypeptides. The individual polypeptides are colored in red and blue. **C.** The same two interacting *Kp*FeoA polypeptides in **B** with one polypeptide displayed as space-filling and the other displayed as ribbon, emphasizing the intercalation of one molecule into the other. **D.** Ribbon representation of the *Kp*FeoA-*Kp*FeoA interaction, with key hydrophobic residues represented as gray balls and sticks. Inset: two Ala residues along β5 (Ala^72^ and Ala^74^) intercalate into a Leu zipper motif comprising 4 Leu residues: Leu^26^ and Leu^29^ (along α1) and Leu^58^ and Leu^60^ (along β4). Leu^29^ becomes displaced from the zipper as it interacts with the Ala residues along β4.

This interaction results in a significant “closing” or “clamping” of the *Kp*FeoA SH3-like fold onto its neighboring polypeptide. We compared our crystal structure to the unpublished NMR structure of *Kp*FeoA (PDB ID 2GCX). Superposition of our structure onto 2GCX chain A shows conservation of the global fold, but there areseveral structural changes resulting in a rmsd of ~2.3 Å over 74 Cα atoms (Fig. 4A). This structural deviation likely explains the failure of the NMR model for MR. The most striking structural differences observed are the closing of the space between the β3-β4 turn and that of the C-terminal side of α1 (Fig. 4A). The two residues anchoring the ends of this region are Leu^29^ and Arg^55^. In the NMR structure, this area is quite open: the distance of Cα Leu^29^ to Cα Arg^55^ is ~10.5 Å. In stark contrast, our crystal structure reveals that this region has closed along β5 of the neighboring molecule: the distance of Cα Leu^29^ to Cα Arg^55^ has decreased to ~6.6 Å, representing a nearly 4 Å narrowing. Previous analyses of the NMR structure of £cFeoA have indicated dynamicism to be present with this same region and, in particular, along β4.^13^ Thus, we propose that 2GCX represents the “open” form of *Kp*FeoA, and that our structure represents the “closed” form of *Kp*FeoA bound to a protein partner.

**Figure 4.**
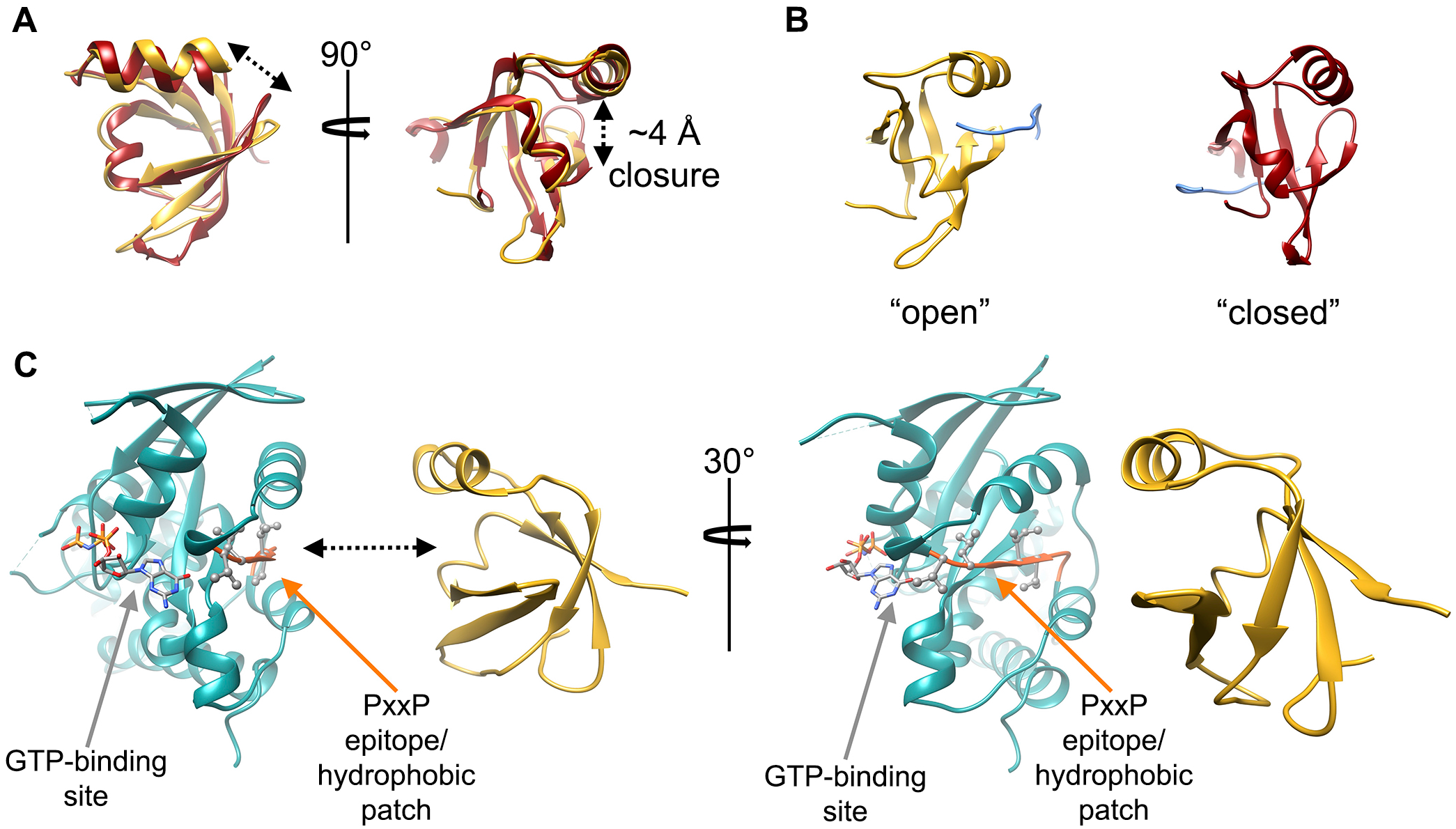
Structural analysis of *Kp*FeoA. **A.** Superposition of our crystal structure of *Kp*FeoA (red) with the NMR structure of *Kp*FeoA (goldenrod; PDB ID 2GCX). While the global fold is conserved, there is a significant closure (~4 Å) of the C-terminal side of α1 and the β3-β 4 turn in our structure compared to the NMR structure. **B.** Docking studies of the 10 amino acid sequence (cornflower) postulated to be the FeoA recognition site along NFeoB. In the “open” NMR structure (goldenrod), the 10-mer peptide docks precisely in the same location we observe interactions between *Kp*FeoA dimers in our crystal structure. In the “closed” crystal structure (red), the 10-mer peptide fails to dock in the “C-shaped” clamp due to the closing of this binding region. **C.** We hypothesize that the open form of *Kp*FeoA (goldenrod) binds to the PxxP recognition site on NFeoB (orange; Leu and Val residues shown as ball and stick of *Kp*NFeoB bound to GMP-PNP, PDB ID: 2WIC). We postulate this binding can be directly communicated to the GTP-binding site through the short helical loop containing hydrogen bonds to the guanine nucleobase. The right panel of **C** represents a 30° rotation of the left panel of **C**.

We believe our structure points to the location along FeoA that may mediate interactions with its corresponding binding site along NFeoB. To test our “open-to-closed” hypothesis, we performed *in silico* docking experiments^21–22^ of both *Kp*FeoA models with a hydrophobic 10 AA sequence (LGCPVIPLVS) representing the postulated partner binding site present on NFeoB. We excised the structure of this 10-mer directly from the crystal structure of *Kp*NFeoB bound to GMP-PNP (PDB ID 2WIC).^23^ Wholly consistent with our hypothesis, the lowest-energy docking model of this peptide with the NMR structure of *Kp*FeoA predicts binding directly within the “C-shaped” clamp of the “open” conformer (Fig. 4B). In contrast and as predicted, the lowest-energy docking model of this peptide with our crystal structure of *Kp*FeoA fails to dock into the now “closed” binding site (Fig. 4B).

In light of these data, we propose a mechanism of FeoA-NFeoB interactions that may link to the status, or alter the state, of bound nucleotide (Fig. 4C). We posit that the “open” conformer of *Kp*FeoA uses its “C-shaped” clamp region (defined by α1 and β4) to interact with the PxxP recognition site on NFeoB. This location is rich in hydrophobic residues (Val and Leu) that would likely intercalate into the Leu zipper along *Kp*FeoA (Fig. 4C), similar to what we observe in our crystal structure (Fig. 3D). Moreover, hydrophobic residues are strongly conserved at, or adjacent to, the Leu zipper location despite low overall conservation of FeoA sequence, emphasizing the functional importance of hydrophobicity in this vicinity. We envision a scenario in which the FeoA-NFeoB binding event is communicated to, or even linked to the state of, GTP/GDP bound on the surface of NFeoB. In the both the GMP-PNP- and GDP-bound *Kp*NFeoB structures, the guanine nucleobase is connected via hydrogen bonding directly to the PxxP binding site by a short helical turn.^23^ Thus, FeoA binding at this site could either increase the rate of GTP hydrolysis, facilitate nucleotide release, or both.

Our discovery of a site for FeoA-mediated protein-protein interactions opens up several exciting avenues for future research on the Feo system. Numerous studies have demonstrated that FeoA is a virtually indispensable component of the bacterial Feo system,^24–27^ likely due to FeoA’s regulatory role in modulating ferrous iron import through its interaction with FeoB. Furthermore, the disruption of protein-protein interactions mediated by complex surface recognition sites along SH3-like domains like FeoA is an active area of pharmacological development in eukaryotes.^20, 28^ This targeted approach could extend to antibiotic development aimed a nutrient uptake pathways such as Feo. We imagine a future scenario in which small, hydrophobic molecules could be developed to disrupt FeoA-FeoB interactions as a novel means of attenuating bacterial virulence through the limitation of iron uptake.

## ASSOCIATED CONTENT

Supporting Information is available: Methods and materials, purification and gel filtration data, additional structural images and tables, and representative partial sequence alignments (PDF)

## ACKNOWLEDGMENT

The authors wish to thank Zdzislaw Wawrzak for his knowledgeable help.

## ABBREVIATIONS

ASU: asymmetric unit
Feo: ferrous iron transport system
GDP: guanosine-5’-diphosphate
GMP-PNP: 5’-guanylyl imidodiphosphate
GTP: guanosine-5’-triphosphate
MR: molecular replacement
NMR: nuclear magnetic resonance
SAD: single-wavelength anomalous dispersion
SH3: Src-homology 3-like fold

---

**SYNOPSIS TOC.**
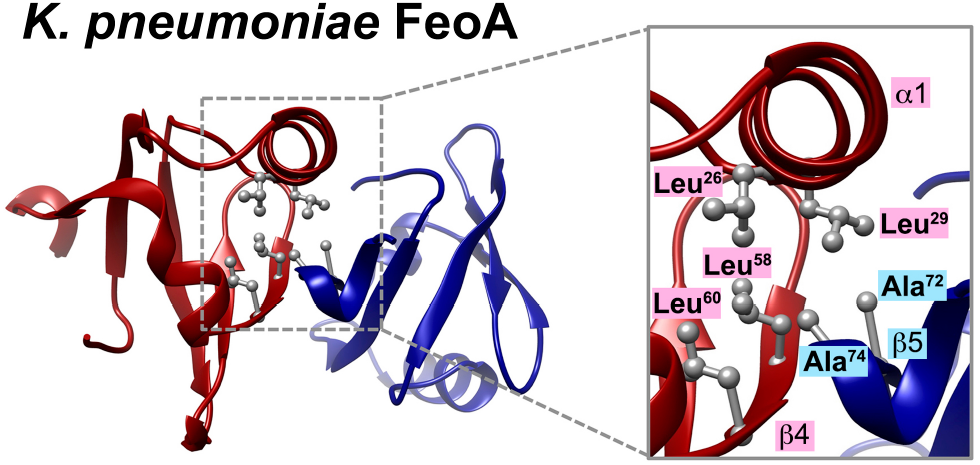
The crystal structure of *K. pneumoniae* FeoA reveals a site for protein-protein interactions mediated by a prokaryotic SH3-like domain. Peptide docking implicates this location as the site of interaction between FeoA and FeoB. These protein-protein interactions are likely utilized to control ferrous iron uptake, a key nutrient acquisition pathway used by pathogenic bacteria to establish infection in acidic and/or anaerobic niches within human hosts. Image for Table of Contents Usage only:

