## Supplemental Information for "The Crystal Structure of Klebsiella pneumoniae FeoA Reveals a Site For Protein-Protein Interactions"

### Supplementary Methods

***Materials.*** The pET-21a(+) expression plasmid was purchased from EMD-Millipore (MilliporeSigma). A modified BL21(DE3) *E. coli* expression cell line in which the gene for the multidrug exporter AcrB (a common contaminant from *E. coli* membranes) had been deleted (BL21(DE3) ΔacrB) was a generous gift of Prof. Edward Yu (Iowa State University). All materials used for buffer preparation, protein expression, and protein purification were purchased from RPI, MilliporeSigma, and/or VWR and were used as received.

***Cloning, Expression, and Purification of KpFeoA.*** DNA encoding for the gene corresponding to FeoA from *Klebsiella pneumoniae* (*subsp. pneumoniae*) (Uniprot identifier A0A0M1TF23) (*Kp*FeoA) was commercially synthesized by GenScript (Piscataway, NJ), with an additionally engineered DNA sequence encoding for a C-terminal TEV-protease cleavage site (ENLYFQS). This gene was subcloned into the pET-21a(+) expression plasmid using the NdeI and XhoI restriction sites, encoding for a C-terminal $(His)_6$ affinity tag when read in-frame. The complete expression plasmid was transformed into chemically competent BL21(DE3) Δ*acrB* cells, spread onto Luria-Bertani (LB) agar plates supplemented with 100 μg/ml ampicillin, and grown overnight at 37°C. Colonies from these plates served as the source of *E. coli* for small-scale starter cultures (generally 100 mL LB supplemented with 100 μg/ml ampicillin). Large-scale expression of *Kp*FeoA was accomplished in 12 baffled flasks each containing 1 L sterile LB supplemented with 100 μg/ml (final) ampicillin and inoculated with a pre-culture. Cells were grown by incubating these flasks at 37°C with shaking of 200 r.p.m. until $OD_{600}$ reached ~0.6-0.8. The flasks containing cells and media were then chilled to 4°C for 2 h, after which protein expression was induced by the addition of isopropyl β-D-l-thiogalactopyranoside (IPTG) to a final concentration of 1 mM. The temperature of the incubator shaker was lowered to 18°C with continued shaking of 200 r.p.m. After ~18-20 h, cells were harvested by centrifugation at 4800×*g*, 10 min, 4°C. Cell pellets were subsequently resuspended in resuspension buffer (50 mM Tris, pH 7.5, 200 mM NaCl, 5% (v/v) glycerol), flash-frozen on $N_{2(l)}$, and stored at -80°C until further use.

For SeMet-substituted *Kp*FeoA, a single colony from LB agar plates supplemented with 100 μg/ml ampicillin (*vide supra*) was used to inoculate 5 mL of sterile LB supplemented with 100 μg/ml ampicillin (final). After ~8 h, ~1 mL of this culture was added to 100 mL of 1x minimal media composed of 1x (final) M9 salts, 0.4% (w/v; final) D-glucose, 2 mM (final) $MgSO_4$, 100 μM (final) $CaCl_2$ supplemented with 100 μg/ml ampicillin. This culture was then grown by incubating these flasks at 37°C with shaking of 200 r.p.m overnight. Large-scale expression of SeMet *Kp*FeoA was accomplished in 12 baffled flasks each containing 1 L sterile 1x supplemented minimal media with 100 μg/ml (final) ampicillin and inoculated with the minimal media pre-culture. Cells were grown by incubating these flasks at 37°C with shaking of 200 r.p.m. until $OD_{600}$ reached ~0.6-0.8. After 2 hr cold shock at 4°C, each 1 L flask of 1x minimal media was supplemented with a solution delivering 100 mg *L*-Phe, 100 mg *L*-Lys, 100 mg *L*-Thr, 50 mg *L*-Ile, 50 mg *L*-Leu, 50 mg *L*-Val, and 60 mg *L*-SeMet. These flasks were then incubated at 18°C with shaking of 200 r.p.m. for 15 min. After this incubation period, protein expression was induced by the addition of IPTG to a final concentration of 1 mM. Protein expression and cell harvest were accomplished as with the native protein (*vide supra*).

All steps for the purification of *Kp*FeoA were performed at 4°C unless otherwise noted. Frozen cells were thawed and stirred until the solution was homogeneous. Solid phenylmethylsulfonyl fluoride (PMSF; ~50-100 mg) was added immediately prior to cellular disruption using a Q700 ultrasonic cell disruptor (QSonica) set to 70% maximal amplitude, 30 s pulse on, 30 s pulse off, for a total pulse on time of 12 min. Cellular debris was cleared by ultracentrifugation at 163000×*g* for 1 h. The supernatant was then applied to a 5 mL HiTrap IMAC FF column (GE Healthcare) that had been charged with $Ni^{2+}$ and equilibrated with 8 column volumes of wash buffer (50 mM Tris, pH 8.0, 200 mM NaCl, 10% (v/v) glycerol) with 21 mM imidazole. The column was then washed with 12 column volumes of wash buffer with 30 mM imidazole. Protein was then eluted by wash buffer containing 300 mM imidazole. Fractions were concentrated using a 15 mL Amicon 3 kDa molecular-weight cutoff (MWCO) spin concentrator (MilliporeSigma). Protein was then applied to a 120 mL Superdex 75 (GE Healthcare) gel filtration column that had been pre-equilibrated with 25 mM Tris, pH 7.5, 200 mM NaCl, and 5% (v/v) glycerol. The eluted fractions of the colorless *Kp*FeoA, which corresponded to either dimeric protein (~20 kDa) or monomeric protein (~10 kDa), were pooled and concentrated with a 4 mL Amicon 3 kDa MWCO spin concentrator. All additional size-exclusion experiments were performed in a similar manner. Protein concentration was determined using the Lowry assay, and purity was assessed via 15% SDS-PAGE analysis.

***Crystallization, Data Analysis, and Structure Determination.*** Crystals of native and SeMet-substituted *Kp*FeoA were obtained by sitting-drop vapor-diffusion using MiTeGen-XtalQuest Plates with a 1:1 (v:v) *Kp*FeoA (~7.5 mg/mL) and reservoir solution mixture at room temperature. The precipitant solution for both native and SeMet *Kp*FeoA consisted of 2.0 M ammonium sulfate, 0.1 M ammonium fluoride, and 3% (v/v) glycerol. Small three-dimensional, colorless octagons appeared within ~24 hr and reached their maximal size within ~2-3 days. Crystals were transferred into cryoprotectant consisting of 1.5 M ammonium sulfate and 50% (v/v) glycerol. After soaking for ~1 min, crystals were then looped, flash-frozen, and stored at 77 K.

Data sets were collected on beamline LS-CAT 21-ID-D at the Advanced Photon Source, Argonne National Laboratory, using a Dectris Eiger 9M detector. Data were processed automatically with Xia2.[1] Phases and initial models of SeMet *Kp*FeoA were generated using the AutoSol and AutoBuild programs in Phenix.[2] This model served as the input for solving native *Kp*FeoA datasets via molecular replacement using Phaser in Phenix.[2] Extended model building and refinement cycles were performed in Coot[3] and REFMAC5[4], respectively. Final validations were performed using Phenix Validate[2], and the final model consists of residues 1-80; the final 10 residues (81-90) comprising chiefly the purification tag are not visible in the electron density map. Data collection and refinement statistics are presented in Table S1. Structural overlays, Cα RMSD values, and surface representations were generated and calculated by utilizing UCSF Chimera (v. 1.11)[5], whereas electron density map overlays were generated using Mac PyMOL (v. 1.7.4.5). Topology images were generated in part by the Pro-Origami server.[6] The atomic coordinates for native *Kp*FeoA have been deposited in the Protein Data Bank (deposition ID: 6E55).

***Docking and Bioinformatics.***

Docking models were created using the ClusPro online server[7-8] with an input of either the *Kp*FeoA crystal structure (chain A; PDB ID 6E55) or the *Kp*FeoA NMR structure (chain A; PDB ID 2GCX) and the 10 amino acid peptide corresponding to LGCPVIPLVS excised from the crystal structure of *Kp*NFeoB bound to GMP-PNP (PDB ID 2WIC)[9]. Docking was performed without modification of the default restraints. The models shown represent those with the lowest balanced, weighted score, but these models are generally representative of observed behavior among all output models.

For bioinformatics analyses, all FeoA sequences were obtained from the Universal Protein Resource (UniProt) Knowledgebase and Reference Clusters (http://www.uniprot.org) or the National Center for Biotechnology Information (http://www.ncbi. nlm.nih.gov/). Multiple sequence alignments were performed using JalView v. 2.7[10] implementing the ClustalW algorithm[11] and the Blosum62 matrix[12].

**Figure S1.** 15% SDS-PAGE analyses of purified *Kp*FeoA. Molecular weight ladders are located at the far left. The middle panel represents the purification of *Kp*FeoA via $Ni^{2+}$ immobilized metal-affinity chromatography (IMAC). Lane 1: the applied soluble fraction; lanes 2 and 3: column flow-through; lane 4: 30 mM imidazole column elution; lane 5: 300 mM imidazole column elution. The right panel represents the purification of *Kp*FeoA via size-exclusion chromatography (SEC) (*vide infra*). Lane 6: dimeric *Kp*FeoA; lane 7: monomeric *Kp*FeoA.

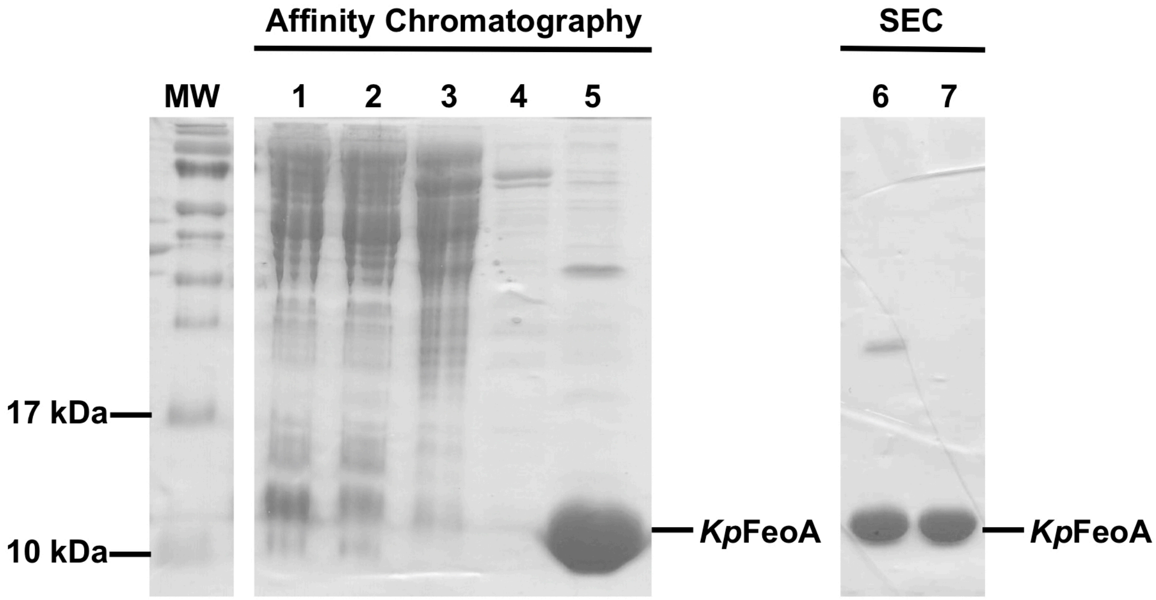

**Figure S2.** Gel filtration data demonstrating static oligomerization of *Kp*FeoA. IMAC-purified *Kp*FeoA exists in two oligomeric states (red trace), whose retention volumes are consistent with dimeric and monomeric *Kp*FeoA. Dimeric *Kp*FeoA was then pooled, diluted, and reinjected over a period of ~6 hr (orange trace). Dimeric *Kp*FeoA is chiefly maintained, indicating a dynamic equilibrium is not present on this time scale.

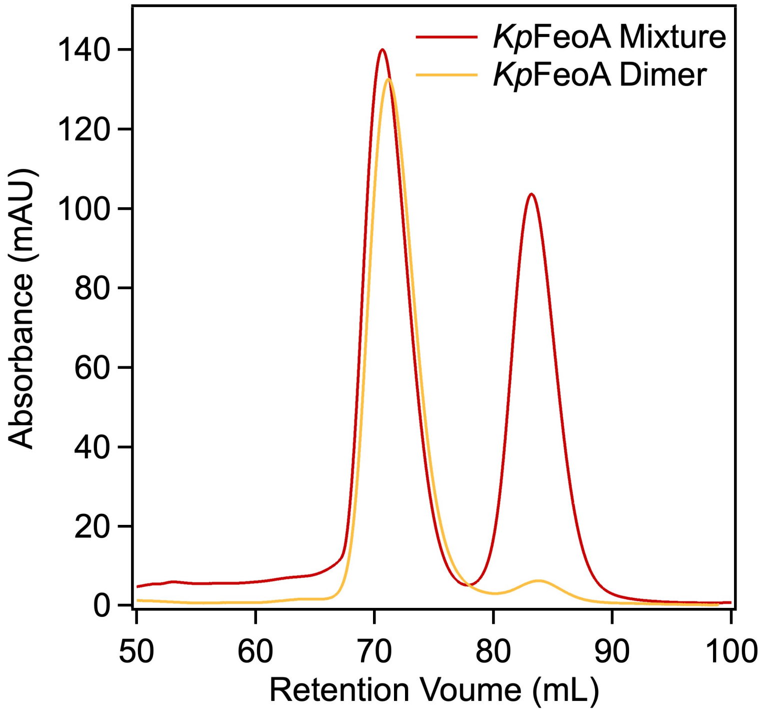

**Figure S3.** Overall structure and electron density map of a single asymmetric unit of *Kp*FeoA. The $2F_o$-$F_c$ electron density map (gray) is contoured at 1σ, and water molecules are explicitly shown.

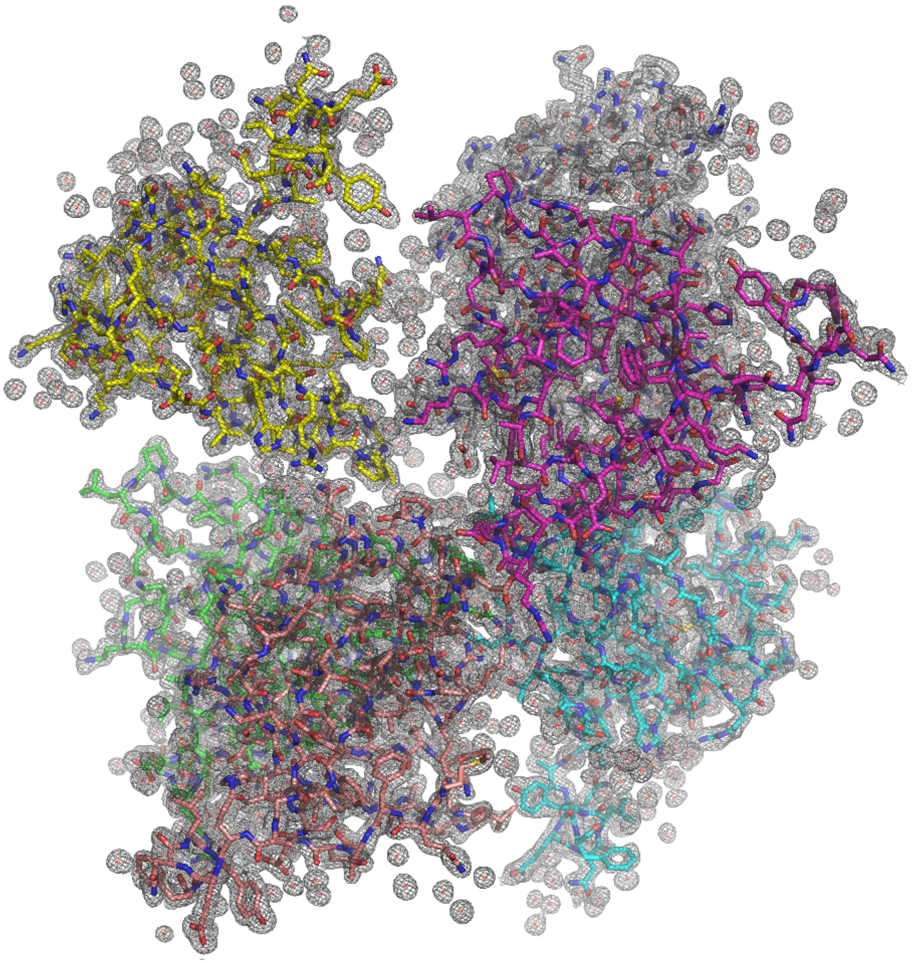

**Figure S4.** Crystal packing and unit cell of *Kp*FeoA showing 5 asymmetric units.

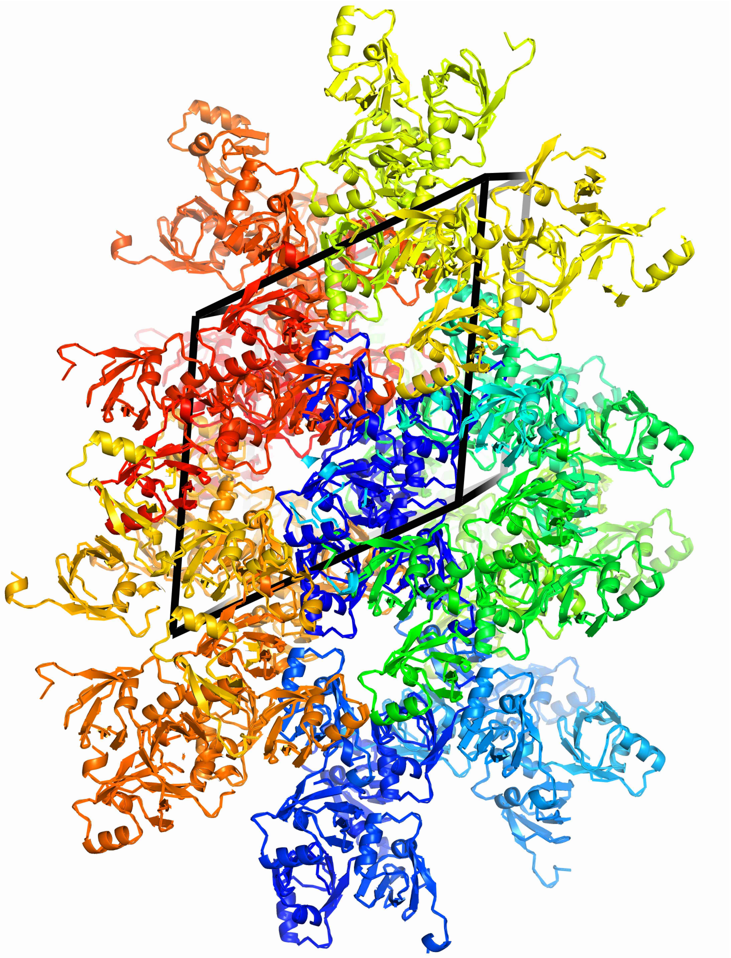

**Table S1.** Data collection and refinement statistics for native and SeMet *Kp*FeoA. Values in parentheses are for the highest resolution shell.

| | ***Kp*FeoA Native** | ***Kp*FeoA SeMet** |
|---|---|---|
| **Data collection** | | |
| Wavelength (Å) | 0.97856 | 0.97849 |
| Space group | $P3_1$ | $P3_121$ |
| Cell dimensions | | |
| *a*, *b*, *c* (Å) | 79.01, 79.01, 74.66 | 45.390, 45.390, 74.050 |
| Resolution (Å) | 36.10 – 1.57 (1.63 – 1.57) | 34.72 – 1.76 (1.79 – 1.76) |
| $R_{merge}$ | 0.067 (1.553) | 0.145 (2.720) |
| I / σ (I) | 15.7 (1.3) | 13.9 (1.1) |
| Completeness (%) | 99.5 (99.5) | 97.1 (89.4) |
| Multiplicity | 7.0 (6.7) | 19.5 (17.3) |
| CC½ (%) | 100 (47) | 100 (52) |
| **Refinement** | | |
| $R_{work}$ / $R_{free}$ | 0.174 / 0.193 | |
| *B*-factors (Å$^2$) | | |
| Protein | 15.73 | |
| Water | 22.35 | |
| R.m.s. deviations | | |
| Bond lengths (Å) | 0.009 | |
| Bond angles (°) | 1.330 | |
| Ramachandran plot (%) | 93.6 / 6.4 / 0.0 | |
